# A topological characterization of DNA sequences based on chaos geometry and persistent homology

**DOI:** 10.1101/2021.01.31.429071

**Authors:** Dong Quan Ngoc Nguyen, Phuong Dong Tan Le, Lin Xing, Lizhen Lin

## Abstract

Methods for analyzing similarities among DNA sequences play a fundamental role in computational biology, and have a variety of applications in public health, and in the field of genetics. In this paper, a novel geometric and topological method for analyzing similarities among DNA sequences is developed, based on persistent homology from algebraic topology, in combination with chaos geometry in 4-dimensional space as a graphical representation of DNA sequences. Our topological framework for DNA similarity analysis is general, alignment-free, and can deal with DNA sequences of various lengths, while proving first-of-the-kind visualization features for visual inspection of DNA sequences directly, based on topological features of point clouds that represent DNA sequences. As an application, we test our methods on three datasets including genome sequences of different types of Hantavirus, Influenza A viruses, and Human Papillomavirus.

## 1 Introduction

The last few decades have witnessed a surge in the growth of methods that are devoted to analyzing the similarities among DNA sequences to obtain the corresponding genetic information. Despite these diverse methods, they can be classified into graphical representation of DNA sequences and other techniques based on numeric representations. Methods based on graphical representation of DNA sequences have contributed significantly to the general area of DNA similarity analysis. One of the main ideas used in graphical representation is to realize each nucleotide base, say A, C, G, T as a point in a Euclidean space, normally, ℝ^*n*^ for some small values *n* between 2 and 5. One then obtains a collection of points in ℝ^*n*^ that represents a DNA sequence. Based on these collections of points, many papers are devoted, in combination with other techniques, such as signal processing (see, for example, [1]), to compare similarities among DNA sequences. The most frequently used similarity measures for analyzing differences or similarities between DNA sequences based their corresponding graphical representation are Euclidean distances or correlation angles. For papers that adopt graphical representation for DNA similarity analysis, the reader is referred to references [2, 3, 4, 5, 6, 7, 8, 9, 10, 11, 12, 13, 14, 15, 16, 17, 18, 19, 20, 21].

Other methods that do not involve graphical representation but instead use numeric representation are based on mapping each nucleotide base to a number, and thus each DNA sequence can be mapped to a number sequence. In order to compare differences or similarities among DNA sequences, other *ad hoc* methods are utilized to compare their corresponding number sequences. The reader is referred to, for example, work [22, 23, 24, 25, 26, 27, 28, 29], and their references therein for these non-graphical representation based methods.

A common theme in graphical representation based methods is to rely on *simply constructed geometric object*, say curves in Euclidean spaces that represent DNA sequences to apply for DNA similarity analysis. Although there were apparent successes in using such methods, the employed techniques pose some technical difficulties: (i) curves constructed via graphical representations may not truly represent the geometry of the corresponding DNA sequences due to degeneracy or self-crossings (ii) these methods may lead to poor performance due to not being able to effectively deal with short and long DNA sequences simultaneously. In an attempt to overcome these technical difficulties, our paper provides a completely novel method for DNA similarity analysis based on a combination of graphical representation with tools from algebraic topology in particular persistent homology. Our method also employs a graphical representation as the first step to transform each DNA sequence into a collection of points in a Euclidean space ℝ^*n*^ which can be viewed as **point clouds** in topological data analysis. Instead of using simple geometry-based methods for analyzing these data points, we apply tools from algebraic topology such as persistent homology to obtain topological signatures of these point clouds that are signified via their persistence diagrams. Each DNA sequence, thus, is in correspondence with its unique persistence diagrams which encodes its topological signatures such as how many connected components, or how many 1-dimensional holes are present in the topology of DNA sequences. Using the well-known Wasserstein distance (which will be reviewed later) for such persistence diagrams, our key observation is that the similarities among DNA sequences are reflected by the Wasserstein distances among their corresponding persistence diagrams. One important feature of our method is its highly distinctive visuality of geometry and topology of DNA sequences. In other words, the DNA sequences, in many cases, can be immediately distinguished by highly distinctive visuality of their corresponding persistence diagrams which are diagrams in ℝ^2^. Another outstanding feature in our method is that it can effectively deal with various lengths of DNA sequences. Our topological framework is general, alignment-free, and can deal with DNA sequences with only partial genome information while providing first-of-the-kind visualization features.

The rest of the paper is organized as follows. In Section 2, we introduce a new higher dimensional (4-dimensional) representation of DNA sequence based on chaos geometry. This new 4-D representation will serve as the basis for building our topological representation of the DNA sequences. In Section 3, we review some basic notions related to persistence homology and formally introduce out method in Section 4. We apply our method for analyzing three datasets in Section 5.

## 2 Chaos 4-dimensional Representation

In this section, we employ 4**-dimensional chaos** (see [30]) to transform a DNA sequence into a finite set of points in ℝ^4^ that can be viewed as a 4-dimensional representation of the DNA sequence. To the best of our knowledge, this is the first time that chaos in higher dimensional space has been adopted to encode DNA sequences although chaos game in 2-dimensional space was already used to represent DNA sequences in previous work (see, for example, [31]). Let *α* be a DNA sequence of length *n* of the form *β*_1_*β*_2_ ⋯ *β*_*n*_, where the *β_i_* denotes one of 4 nucleotide bases A, C, G, T. We first map *β*_1_*β*_2_ ⋯ *β*_*n*_ to a sequence of integers (*a*_1_, *a*_2_, …, *a*_*n*_), where

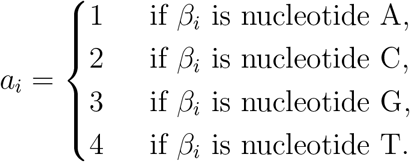

Set *e*_1_ = (1, 0, 0, 0), *e*_2_ = (0, 1, 0, 0), *e*_3_ = (0, 0, 1, 0), and *e*_4_ = (0, 0, 0, 1), the four standard unit vectors in ℝ^4^. Using the 4-dimensional chaos, we construct the finite set of points *X_α_* in ℝ^4^, consisting of points *b*_1_, …, *b_n_* ∈ ℝ^4^ as follows.

i. 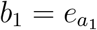; and
ii. for each 2 ≤ *k* ≤ *n*, set 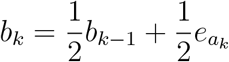.

Intuitively, *b*_*k*_ is the *k*-th point in the 3-simplex that is chosen to be the midpoint of the line segment that connects the (*k* − 1)-th point and the vertex 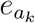. The map that transforms each DNA sequence *α* = *β*_1_*β*_2_ ⋯ *β*_*n*_ to a finite set of points *X_α_* = {*b*_1_, …, *b_n_*} in ℝ^4^, is called the **Chaos** 4**-dimensional Representation** (C4DR).

## 3 Persistent homology and persistence diagrams

In this section, we recall the notion of persistent homology and persistence diagrams that we be used in our method for analyzing DNA sequences. One of the main references for persistent homology is [32].

### 3.1 Homology groups of a simplicial complex

#### Definition 3.1.

(*k*-simplex)

Let *k* be a nonnegative integer, and let *u*_0_, …, *u*_*k*_ be *k* +1 points in ℝ^*k*+1^. A *k*-simplex *σ* generated by {*u*_0_, …, *u_k_*} is the convex hull of {*u*_0_, …, *u_k_*}, i.e., the set consisting of all convex combinations of these points that is given by

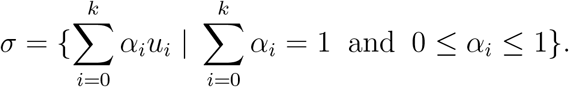

Throughout this paper, we denote by [*u*_0_, …, *u_k_*] the *k*-simplex generated by the points *u*_0_, …, *u_k_*.

Intuitively, a 0-simplex is a point in ℝ, a 1-simplex is a line in ℝ^2^, and a 2-simplex is a triangle in ℝ^3^ (see Figure for illustration of these simplices).

The convex hull of any subset of {*u*_0_, …, *u_k_*} with *d* + 1 elements is also a *d*-simplex, and is called a **face of** *σ*.

#### Definition 3.2.

(simplicial complex)

A simplicial complex Δ is a collection of simplices such that whenever *σ* is a simplex in Δ, all the faces of *σ* are contained in Δ.

A key notion for constructing persistent homology of a set of points in ℝ^*n*^ is a formal sum of *j*-simplices. A **formal sum of** *j***-simplices** is an object of the form ∑_*h*_ *a_h_σ_h_*, where the *a_h_* are real numbers in ℝ such that all but finitely many *a_h_* are zero, and the *σ_h_* are *j*-simplices. Two formal sums ∑_*h*_ *a_h_σ_h_* and ∑_*h*_ *b_h_σ_h_* can be added in a natural way as

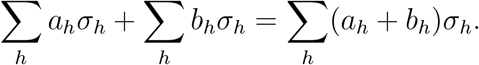

Equipped with this natural addition “+”, the collection of all formal sums ∑_*h*∈*H*_ *a_h_σ_h_* becomes a **group**–a mathematical structure in Algebra in which one can add, and subtract its elements in a similar way as the real numbers do.

Let Δ be a simplicial complex in ℝ^*n*^. For *j* ≥ 0, the *j***-th chain group** 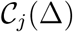 is the **group of all formal sums of** *j***-simplices** ∑_*h*_ *a_h_σ_h_*, where the *σ_h_* are *j*-simplices in Δ. Each formal sum in 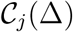 is a *j***-chain**.

There is a natural map *∂_j_* which can send an element in 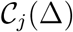 to 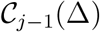 by removing one point from the generating set of a formal sum, and taking the alternating sum of them. It suffices to give the equation of *∂_j_* for each *j*-simplex [*u*_0_, …, *u_j_*] in Δ that is given by

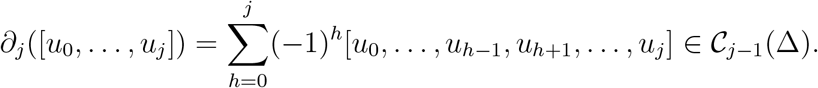

Thus one obtains the map, called the *j***-th boundary map**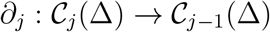.

The *j*-th chain group 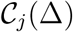 contains two important subgroups 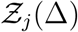 and 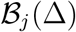. The former, 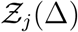, called the *j***-th cycle group**, consists of all *j*-chains *σ* in 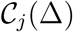 such that *∂_j_* (*σ*) = 0. For example, for any three distinct points *u*_0_*, u*_1_*, u*_2_ in ℝ^2^, the 1-chain *σ* = [*u*_0_*, u*_1_] + [*u*_1_*, u*_2_] + [*u*_2_*, u*_3_] is a 1-cycle since *∂*_1_(*σ*) = 0. The latter, 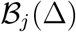, called the *j***-th boundary group**, consists of all *j*-chains *∂_j_*_+1_(*σ*), where *σ* varies over the (*j* + 1)-th chain group 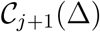. The fundamental theorem of homology implies that 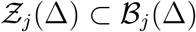.

The following is one of the most important notions that we will use in our method.

#### Definition 3.3.

(homology groups)

Let Δ be a simplicial complex in ℝ^*n*^. For each *j* ≥ 0, the *j***-th homology group** of Δ is the quotient group 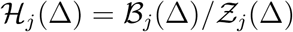.

### 3.2 Persistent Homology Groups and Persistence Diagrams

In this subsection, we recall the notion of persistent homology groups generated by a finite set of points in ℝ^*n*^. Let *X* be a finite set of points, say *u*_1_, …, *u_d_* in ℝ^*n*^, and let *d* denote the standard Euclidean distance in ℝ^*n*^. For any *α* ≥ 0, let 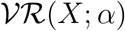 be the collection of all subsets *σ* of *X* such that the Euclidean distance between any two elements in *σ* is at most *α*, that is,

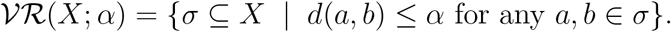

It is easy to verify that 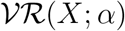 is a simplicial complex called the *α***-Vietoris-Rips complex of** *X*, and thus one can construct homology groups 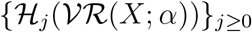 as in Subsection 3.1.

Now take an increasing sequence of nonnegative real numbers, say {*α_i_*}_*i*≥1_ with *α_i_ < α_i_*_+1_ for all *i*. Set *α*_0_ = −∞. For each *i* ≥ 1, set

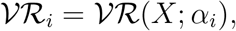

and 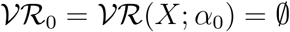. Then one obtains a filtration of the form

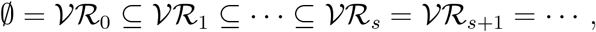

where *α_s_* is large enough such that all pairs of points in *X* are within *α_s_*. For each 0 ≤ *p* ≤ *q* ≤ *s*, there is a natural map 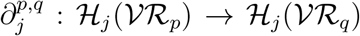 induced from the embedding 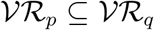. We denote by 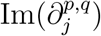 the image of the 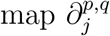.

#### Definition 3.4.

(persistent homology groups)

For each *j* ≥ 0, the *j***-th persistent homology groups of** *X* are the images 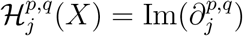 for 0 ≤ *p* ≤ *q* ≤ *s*.

By a *j***-topological feature of** *X*, we mean an element *γ* in 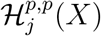 for some 0 ≤ *p* ≤ *s*. The *j***-th persistence diagram of** *X* is a set of points {*b, d*) |0 ≤ *b* < *d*}, where each point (*b, d*) signifies the birth and death times of a *j*-topological feature *γ* of *X*, i.e., *b* is the radius in which *γ* first appears in 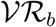 and *d* is the radius in which *γ* gets filled in with a lower dimensional simplex. We denote by 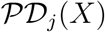 the *j*-th persistence diagram of *X*. In our methods, it suffices to consider only the 0-th and 1-th persistence diagrams, which correspond to topological features of connectedness and 1-dimensional holes of *X*, respectively.

Let *X, Y* be two finite sets of points in ℝ^*n*^. In order to compare topological features of *X, Y* in our methods, we consider the **Wasserstein distance of degree** 1 between 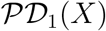 and 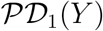, i.e.,

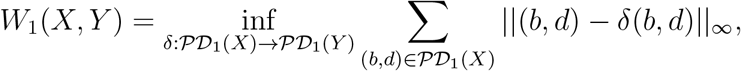

where || · ||_∞_ denotes the *L*_∞_-distance between two points in ℝ^2^.

## 4 Method

Our proposed method for reconstructing a phylogenetic tree of DNA sequences is described in the following algorithm:

(0) (Input) A collection of *n* DNA sequences *α*_1_, …, *α_n_*.
(1) Construct Chaos 4-dimensional Representation (C4DR) of each DNA sequence *α_i_* to obtain a finite set of points 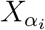 in ℝ^4^ (see Section 2).
(2) Compute the 1st persistence diagrams of the 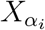 to obtain the sets 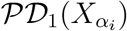 in ℝ^2^, using the notions in Section 3 ^1^.
(3) Compute the distance matrix of dimensions *n* × *n* whose (*i, j*)-entry is the Wasserstein distance 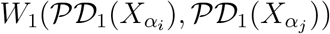.
(4) (Ouput) Construct the phylogenetic tree of the DNA sequences from the distance matrix in Step 3, using UPGMA algorithm (see [33]).

## 5 Results

In this section, we apply out method described in Section 4 to analyzing three datasets: Human Papillomavirus (HPV), Hantavirus, and Influenza A virus.

### 5.1 Human Papillomavirus (HPV)

Let us begin with the dataset of HPV. Human Papillomavirus is mostly responsible for cervical cancer which is the second most common cancer among women (see [34]). We apply our method on the data set of 400 HPV genomes that consist of 12 genotypes 6, 11, 16, 18, 31, 33, 35, 45, 52, 53, 58, and 66. Note that among these genotypes, there are low risk HPV types such as 6 and 11, and high risk HPV types such as 16 and 18. These high risk HPV types are responsible for about 70% of cervical cancer (see [35]). Thus it is an important problem to accurately classify HPV into low and high risk types. In addition, an ideal method should be able to identify HPV genotypes when only partial genomes are available. Our proposed method in Section 4 have all features to become a suitable and good candidate for efficiently classifying HPV genotypes. In addition it has a highly distinctive visuality that can clearly distinguish HPV genotypes in terms of visualization. For example, Figure 1 illustrates identical persistence diagrams of subtypes 11 and 15 of the same HPV genotype 6 whose highly identical visualization shows that they should belong in the same HPV genotype. On the other hand, the distinctive visualization between subtype 44 of HPV genotype 16 and subtype 18 of HPV genotype 18 in Figure 2 shows that they should belong in different genotypes of HPV. In fact the Wasserstein distance between them is 1.548–the maximum distance between any two genome sequences of HPV from the data set.

**Figure 1:**
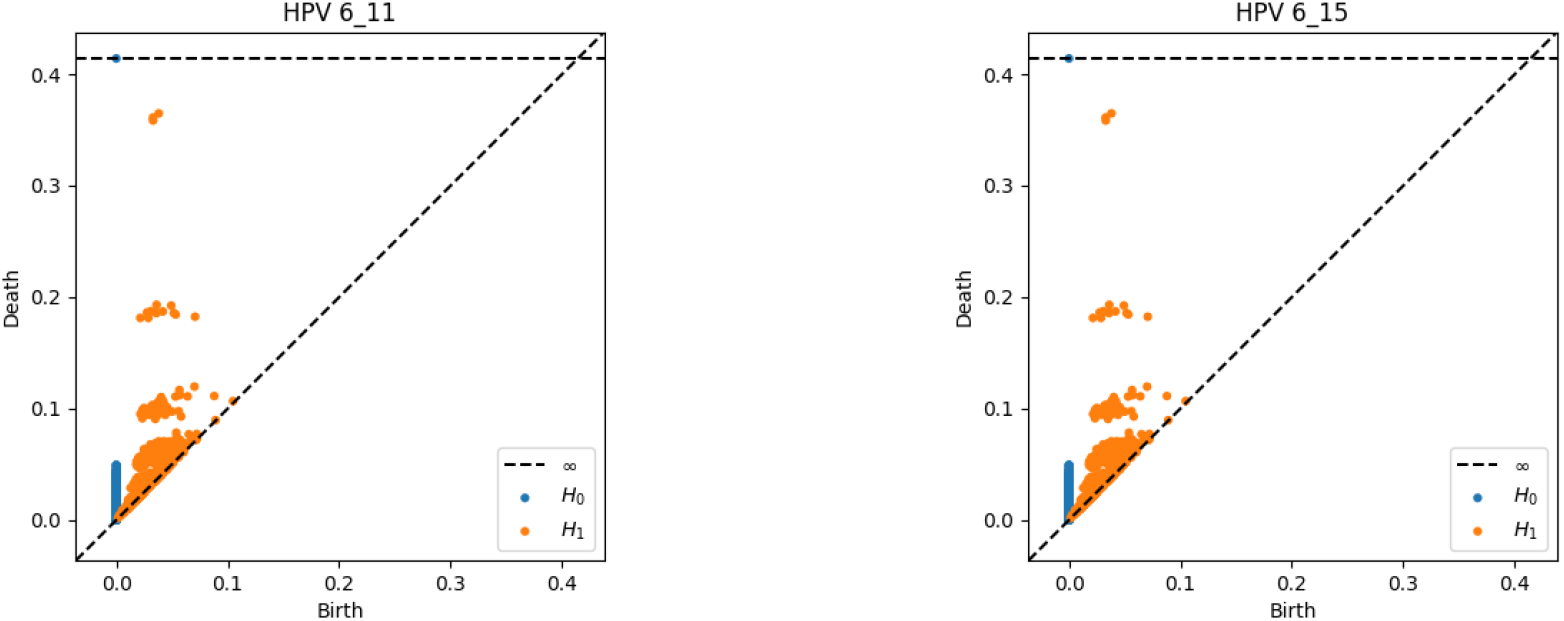
Persistence diagrams of HPV genomes with the minimum Wasserstein distance 0.

**Figure 2:**
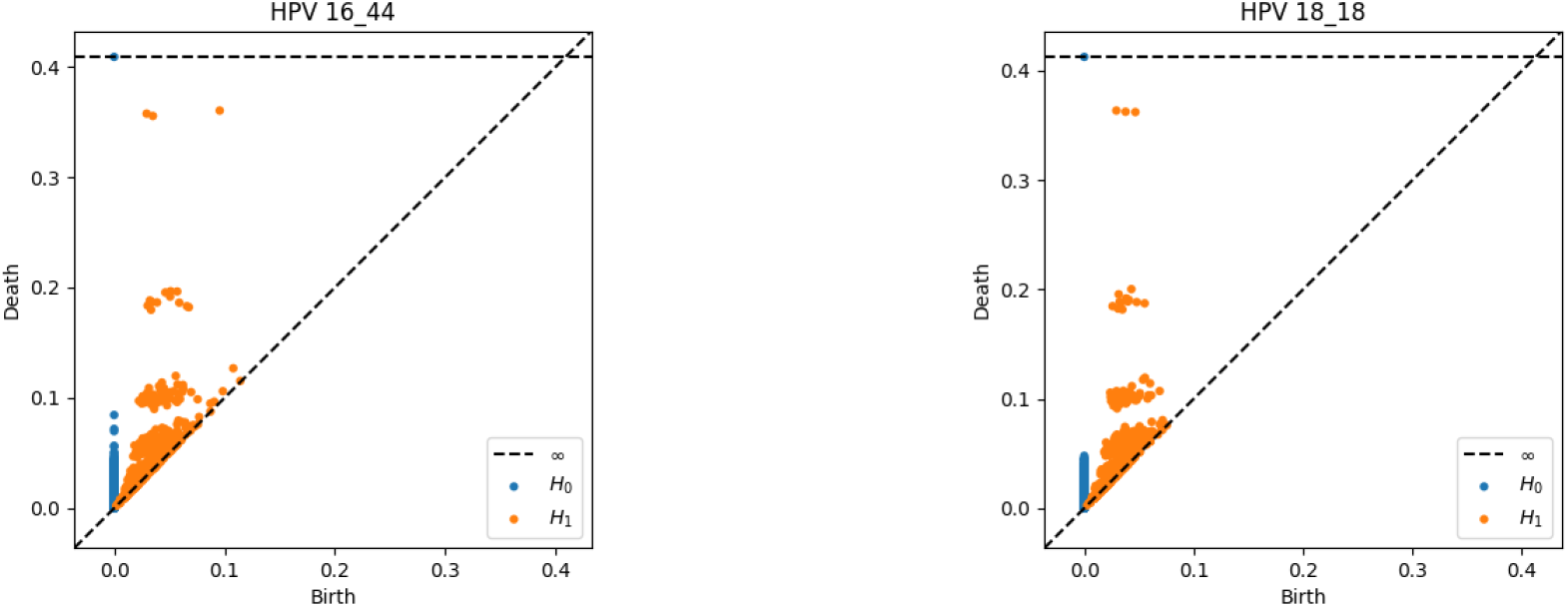
Persistence diagrams of HPV genomes with the maximum Wasserstein distance 1.548.

From the phylogenetic tree of HPV genomes based on our method in Figure 5, our method accurately classifies these HPV genomes into their corresponding genotypes. Note that in [1], their CGR method misplaced one genome of HPV type 11, and they further showed that Clustal Omega (see [36]) is able to classify the dataset of HPV correctly.

### 5.2 Hantavirus (HV)

Hantavirus, named after Hanta River in South Korea, is a recently discovered RNA virus in the family *Bunyaviridae*. HV may infect humans, and some strains of HV can cause possibly fatal diseases in humans such as *Hantavirus hemorrhagic fever with renal syndrome* (HFRS) or *hantavirus hemorrhagic fever with renal syndrome* (HPS). In Eastern Asia, the type of HV that causes HFRS, mainly include Hantaan (HTN) and Seoul (SEO) viruses. In Western European, Russia, and Northeastern China, Puumala (PUU) is the type of HV that causes HFRS. There is another genus of the family *Bunyaviridae* that is called Phlebovirus (PV). We apply our method on the data set of 34 HV genome sequences consisting of 4 different types HTN, SEO, PUU, and PV. The name of these strains are included in the phylogenetic tree (see Figure 3). From Figure 3, we find that our method accurately cluster CGRN strains together whose host is *Rattus norvegicus*. Similarly the method correctly clusters four CGAa strains together whose host is *Apodemus agrarius*. In addition, two CGHu strains whose host is Homo sapiens, is also grouped together.

**Figure 3:**
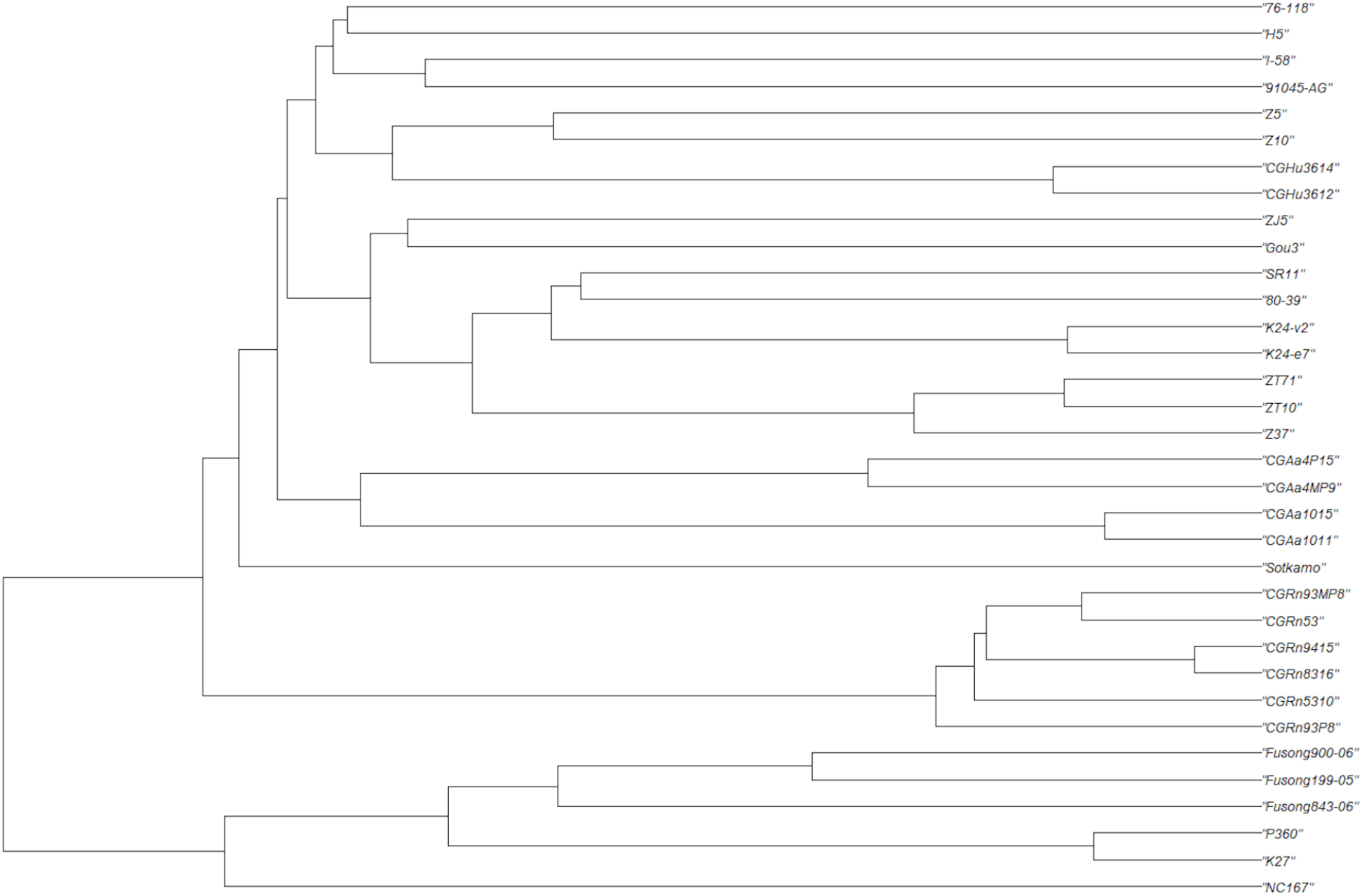
Phylogenetic tree of Hantavirus genomes

Except strain Sotkamo (type PUU) forms an independent leave, we find that all leaves are classified accurately into their corresponding types. But branches of the phylogenetic tree constructed by our method contain both types as sub-branches which are quite different from the results in [37, 38, 39].

### 5.3 Influenza

Influenza A viruses are very dangerous because their hosts have a wide range including birds, horses, swine, and humans. These viruses have been a serious health threat to humans and animals (see [40]), are known to have high degree of genetic and antigenic variability (see [41, 42]). Some subtypes of Influenza A viruses are very lethal including H1N1, H2N2, H5N1, H7N3, and H7N9. We apply our method on the dataset consisting of 38 Influenza A virus genomes. From Figure 4, we find that except A/emperor goose/Alaska/44297-260/2007 (H2N2) and A/turkey/VA/505477-18/2007 (H5N1), the *tips (or leaves)* of the phylogenetic tree of segment 6 of Influenza A virus genomes are clustered correctly into their types, but, for example, for type H1N1 genomes, our method clusters them into 3 distinct subbranches of the tree. We will address this problem in an ongoing work.

**Figure 4:**
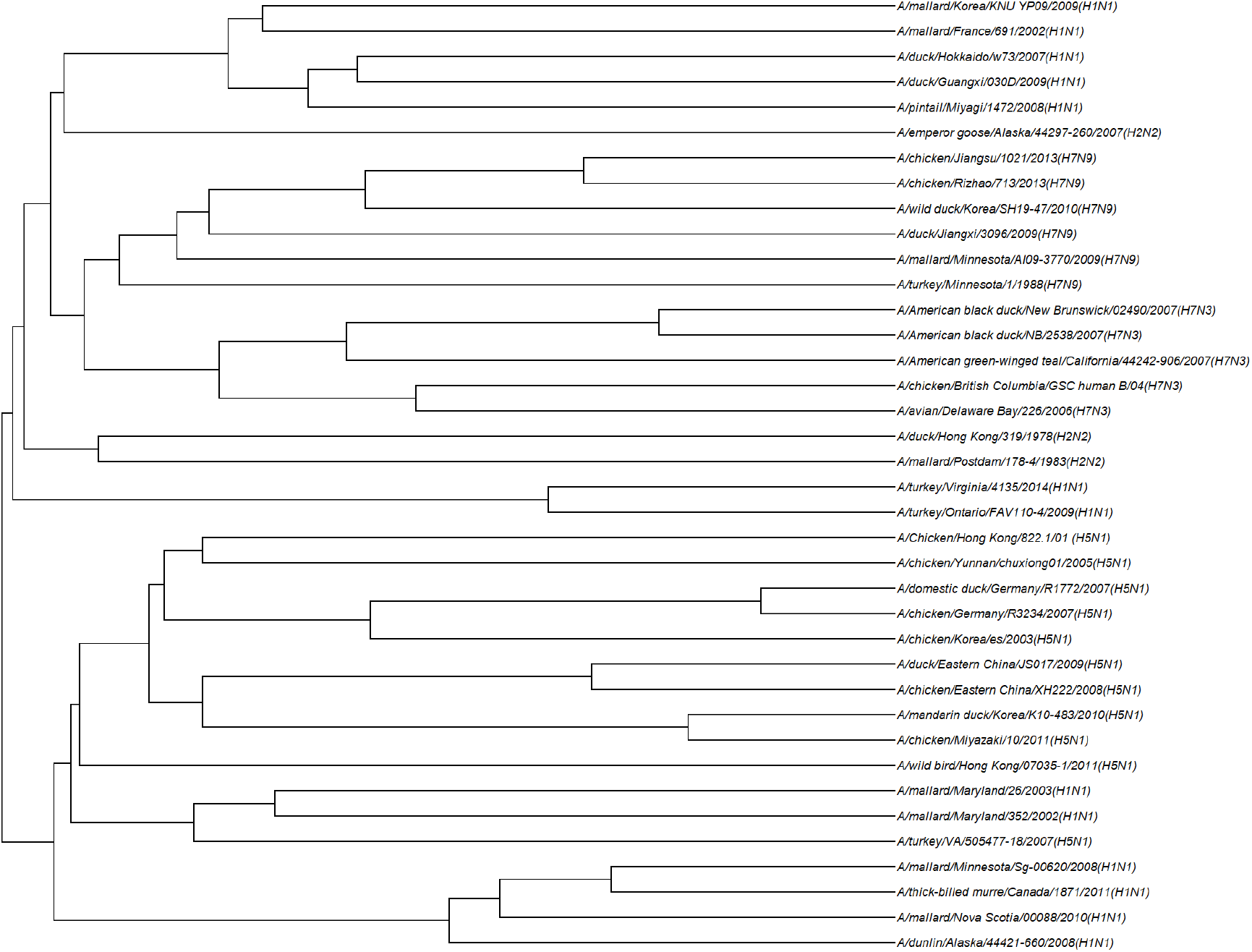
Phylogenetic tree of 38 Influenza A virus genomes

**Figure 5:**
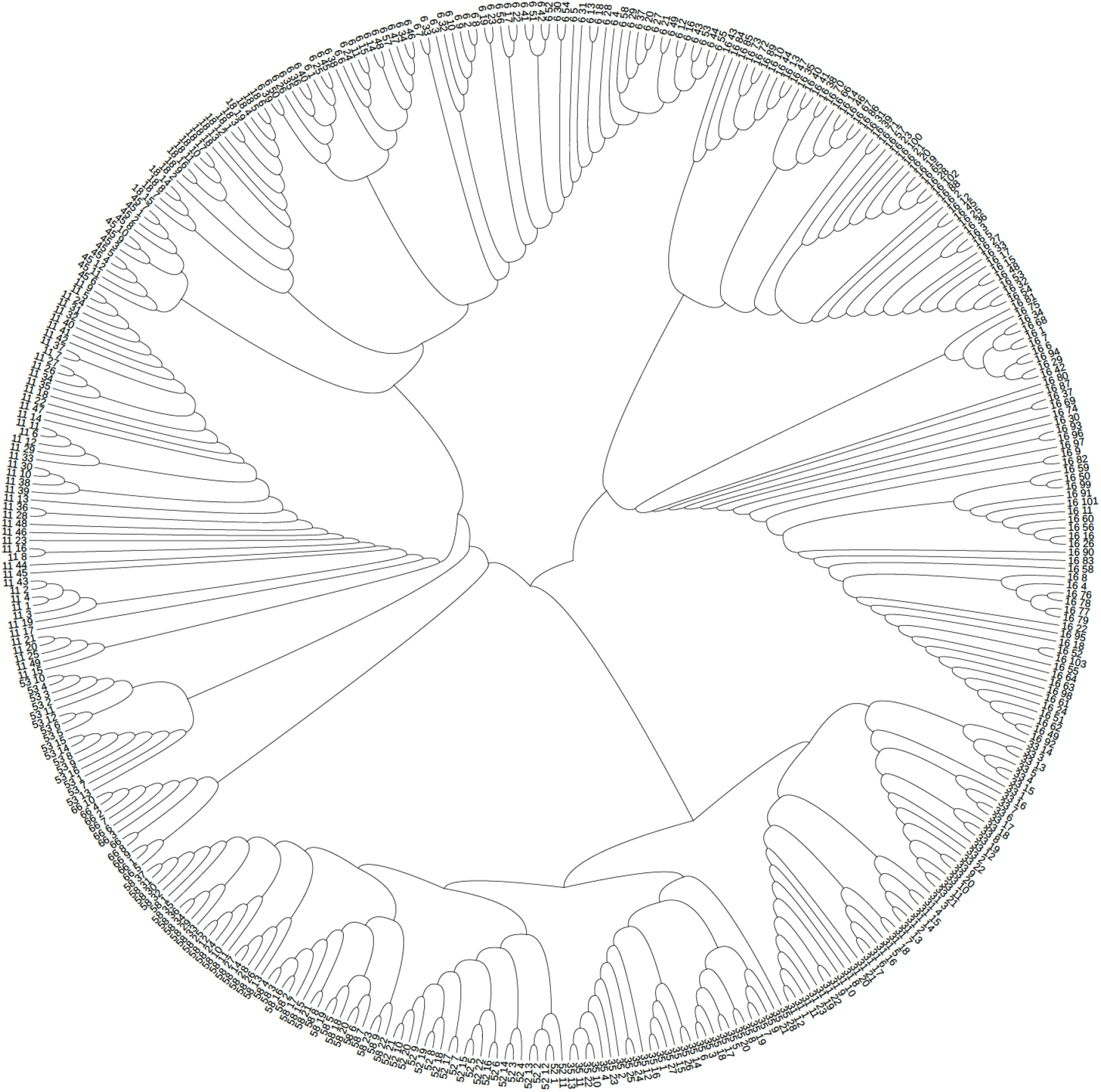
Phylogenetic tree of 400 HPV genomes of 12 genotypes

## Footnotes

1 We use Python packages from https://pypi.org/project/persim/ to compute persistence diagrams and Wasserstein distances.

## Notes

### Competing Interest Statement

The authors have declared no competing interest.

